# Habitat degradation destabilizes complex metacommunities by disrupting dispersal

**DOI:** 10.64898/2026.11.01.763431

**Authors:** Yongzheng Sun, Xiangxin Yin, Zhen Jin, Shirui Bian, Wei Lin, Vit Piskovsky, Philip K. Maini

**Author notes:** Corresponding author: **Correspondence** and requests for materials should be addressed to (Y.S.), (W.L.), (V. P.), (P.K.M.).

## Abstract

Habitat loss and fragmentation are rapidly reshaping ecosystems worldwide, yet their consequences for ecological stability remain unresolved. Classical theory, pioneered by Robert May, established rigorous criteria for stability; however, these apply to spatially well-mixed communities, limiting their relevance to habitat degradation. Here, we develop a metacommunity framework that integrates classical stability theory with spatial dynamics to determine how anthropogenic habitat degradation reshapes ecosystem stability. We show that metacommunities exhibit three distinct dynamical regimes following a perturbation (e.g., drought or forest fire): stable regime (perturbations decay and species persist), reactive regime (perturbations are transiently amplified, which increases extinction risk under repeated disturbances), and unstable regime (perturbations grow, causing extinctions even after a single disturbance). Dispersal fundamentally reshapes these stability transitions, converting unstable communities into reactive ones and reactive communities into stable ones. While homogeneous dispersal progressively stabilizes communities, heterogeneous dispersal maximizes stability at intermediate strengths. We further show that habitat loss and fragmentation can be comparably effective at driving species extinctions by shifting metacommunities from stable to reactive and ultimately unstable regimes, demonstrating that fragmentation *per se* can be as important as habitat loss. Our findings reveal that reactivity is both an early-warning signal of impending instability and a state of elevated extinction risk under recurrent disturbances. By linking stability transitions to changes in population persistence under nonlinear dynamics, our results bridge classical stability theory with spatial ecology and provide a general framework for predicting how ecological complexity and dispersal jointly govern both ecosystem stability and population persistence under habitat degradation.

---

Habitat loss and fragmentation are the dominant threats to ecosystems worldwide, placing many species at risk of extinction [1–4]. By disrupting species dispersal and limiting their access to essential resources, habitat degradation isolates communities within increasingly small and fragmented patches [5–7]. Moreover, these fragmented communities (metacommunities) are increasingly exposed to stochastic perturbations, such as floods or fires, which are intensifying under climate change [8–10]. This raises a central question: how do metacommunities respond to such perturbations – do they recover or collapse?

This question has a long tradition in theoretical ecology, where the ability of communities to recover from perturbations is termed *stability*. In his foundational work, Robert May developed a mathematical framework to assess the linear stability of large ecological communities (i.e., their ability to recover from small perturbations) under random and independent interactions between different members of the community [11]. Subsequent extensions have incorporated features such as interaction diversity [12–14], time delay [15, 16], trophic structure [17], modularity [18, 19], and higher-order interactions [20]. However, these approaches have largely focused on well-mixed communities without explicit spatial structure. Only recently have theoretical frameworks begun to address spatially structured systems (metacommunities) [18, 19, 21–26], yet these remain predominantly abstract and have not been used to examine the effects of habitat loss and fragmentation.

Moreover, while stability determines the final long-term outcome, that is, whether the community recovers from a perturbation or not, it is also important to understand how stable communities recover. Do they directly return back to their original state or is the perturbation transiently amplified? In theoretical ecology, the transient amplification of perturbations is termed *reactivity* [27], which is a key property of complex systems [28–31] and which has been linked to an increased extinction risk if the stochastic perturbations have negative impacts and are common [32], making reactivity particularly relevant as climate change increases the frequency and intensity of ecological disturbances [8]. However, similar to stability, the study of reactivity has been limited exclusively to well-mixed communities and it remains unclear how reactivity emerges in spatially structured metacommunities and how it responds to habitat degradation.

Here, we develop a theoretical framework to investigate the stability and reactivity of spatially structured metacommunities under habitat degradation. Using linear stability analysis and random matrix theory, we examine how habitat loss, fragmentation and dispersal jointly shape the response of ecosystems to perturbations. By linking stability transitions to species extinction under single and recurrent disturbances, our framework connects classical stability theory with population persistence, providing a general basis for understanding how anthropogenic habitat change translates dynamical vulnerability into biodiversity loss.

## Results

### Model overview

Local communities residing in distinct habitats tend to be interconnected as fluxes of individuals, energy, and materials, thereby constituting a metacommunity, as illustrated in Fig. 1a. Following previous work [11, 33], we model a metacommunity composed of *n* interacting species across *m* habitats as a continuous-time dynamical system

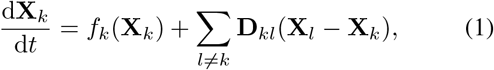

where 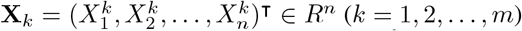 represents an *n*-dimensional vector and 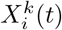 denotes the abundance of species *i* in patch *k* at time *t*. The function *f*_*k*_: *R*^*n*^ →*R*^*n*^ models the network structure and interspecies interactions within patch *k*, with elements 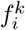 corresponding to common ecological network dynamics, while **D**_*kl*_(**X**_*l*_ − **X**_*k*_) represents dispersal between the same species in different patches.

**FIG. 1.**
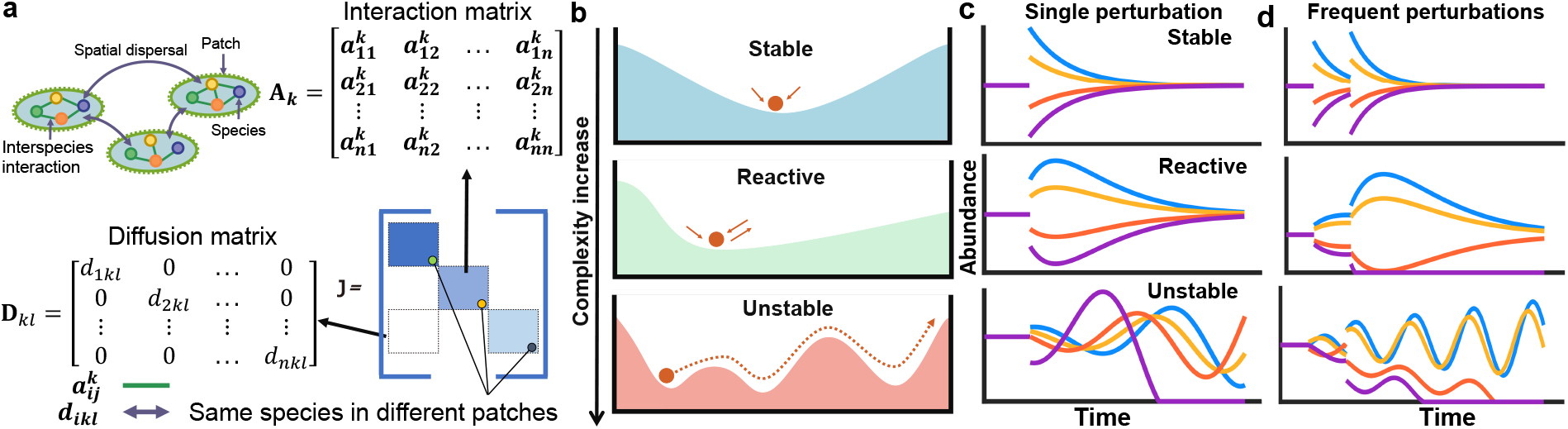
Graphical representation of metacommunities and stability transitions. **a**. Conceptual diagram of a metacommunity. The diagonal blocks of the Jacobian matrix **J** consist of random matrices **A**_*k*_ capturing species-species interactions, while the off-diagonal blocks are diagonal matrices **D**_*kl*_ representing dispersal between patches. **b**. Increasing complexity drives a transition from stability to reactivity and, ultimately, to instability. **c-d**. Schematic responses of species abundances to a single perturbation (**c**) and recurrent perturbations (**d**). Stable metacommunities recover from perturbations and maintain species persistence; reactive metacommunities remain asymptotically stable but transiently amplify perturbations, increasing extinction risk when perturbations recur; unstable metacommunities fail to recover and can experience species extinctions even after a single perturbation.

Here, we measure ecosystem stability (i.e., the ability of a system to return to equilibrium after external perturbations) by local asymptotic stability. For the metacommunity described by Eq. (1), we assume the existence of a feasible equilibrium 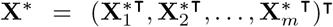 (where all components of **X**^*\**^ are positive) and linearize the dynamics around **X**^*\**^ to assess its local stability. The resulting linearized system can be written as

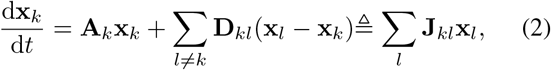

where 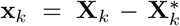 denotes the deviation from the feasible equilibrium. Here, the matrix **A**_*k*_ consists of the elements 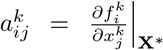 that represent the ecological interaction between species *j* and *i* in patch *k*, where 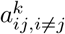 is drawn at random with probability *c* from a fixed probability distribution with standard deviation *σ*, in line with the previous work [11]. Specifically, the off-diagonal elements 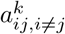 represent the effect of species *j* on species *i* and the diagonal elements 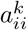 describe the self-regulation of species *i*. The diagonal matrix **D**_*kl*_ defines the dispersal movement between patches *k* and *l*, quantifying the impact of spatial dispersal. For notational simplicity, we combine the within-patch ecological interactions **A** and the between-patch dispersal **D** into a single matrix **J**, known as the Jacobian matrix, illustrated in Fig. 1a.

For metacommunities, stability can be determined by examining the maximum real part of the eigenvalues of matrix **J**, denoted as max ℜ(*λ*(**J**)), which effectively determines whether the system returns to equilibrium after perturbation, as well as its return rate. Specifically, if max ℜ(*λ*(**J**)) *<* 0, then the metacomunity returns to its original state with a positive rate and the system is stable [11]. In contrast, if max ℜ(*λ*(**J**)) *>* 0, the metacommunity never returns to its original state and the system is unstable. Stability characterizes the long-term response to small perturbations. However, certain perturbations around a stable equilibrium may exhibit a transient amplification known as reactivity. Here, we adopt the conventional definition of reactivity, which measures the maximum instantaneous amplification of perturbations [27]

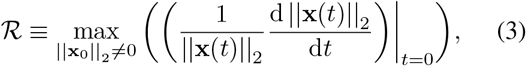

where **x**_0_ is the vector of perturbed species abundances in the system and ||·||_2_ denotes the (Euclidean) magnitude of this perturbation. Mathematically, *ℛ*can be calculated as the largest eigenvalue *λ*_max_(**H**) of the matrix **H** = (**J**+**J**^⊺^)*/*2, i.e., the symmetric part of the Jacobian matrix **J**.

The condition *ℛ>* 0 indicates that the system is reactive, meaning that there exist certain initial perturbations that can be amplified over time. Conversely, if *ℛ<* 0, then all initial perturbations decay over time, and the system is non-reactive. Therefore, the key to assessing the stability and reactivity of metacommunities lies in determining max ℜ(*λ*(**J**)) and *λ*_max_(**H**), respectively. It is worth noting that non-reactive systems are always stable [27, 28, 32], which simplifies the following classification of stability.

In summary, we classify metacommunities into three categories (Fig. 1b): (i) *stable*, if they exhibit stability without reactivity (*λ*_max_(**H**) *<* 0); (ii) *reactive*, if they are stable but exhibit transient amplification (max ℜ (*λ*(**J**)) *<* 0 and *λ*_max_(**H**) *>* 0); and (iii) *unstable*, if they are both reactive and unstable (max ℜ(*λ*(**J**)) *>* 0). Using this classification, we analytically predict the stability and reactivity of metacommunities with diverse interaction types, including random (± */*±), predator–prey (+*/*−), competitive (− */*−), mutualistic (+*/*+), and their combinations listed in Table 1 (see Methods and Supplementary Information for detailed derivation). For simplicity, we assume here that all species share the same self-regulation coefficient *s* and dispersal rate *d*. However, we will also later address the case of heterogeneous dispersal numerically.

**TABLE 1.** Stability and reactivity criteria for metacommunities.

|  | **Random** | **Predator-prey** | **Competition** | **Mutualism** |
|---|---|---|---|---|
| Stability ($d \ll 1$) | $\sigma\sqrt{cn} < s + m'd$ | $\sqrt{cn\sigma^2}\frac{\sigma_1^2}{\sigma^2} < s + m'd$ | $\sqrt{cn\sigma_2^2}\frac{\sigma_3^2}{\sigma_2^2} + c\mathbb{E}(\|z\|) < s + m'd$ | $c(n-1)\mathbb{E}(\|z\|) < s + m'd$ |
| Reactivity ($d \ll 1$) | $\sigma\sqrt{2cn} > s + m'd$ | $\sqrt{2cn\sigma_1^2} > s + m'd$ | $\sqrt{2cn\sigma_3^2} + c\mathbb{E}(\|z\|) > s + m'd$ | $c(n-1)\mathbb{E}(\|z\|) > s + m'd$ |
| Stability ($d \gg 1$) | $\sigma\sqrt{\frac{cn}{m}} < s$ | $\sqrt{\frac{cn\sigma^2}{m}}\frac{\sigma_1^2}{\sigma^2} < s$ | $\sqrt{\frac{cn\sigma_2^2}{m}}\frac{\sigma_3^2}{\sigma_2^2} + c\mathbb{E}(\|z\|) < s$ | $c(n-1)\mathbb{E}(\|z\|) < s$ |
| Reactivity ($d \gg 1$) | $\sigma\sqrt{\frac{2cn}{m}} > s$ | $\sqrt{\frac{2cn\sigma_1^2}{m}} > s$ | $\sqrt{2cn\sigma_3^2} + c\mathbb{E}(\|z\|) > s + d$ | $c(n-1)\mathbb{E}(\|z\|) > s$ |

The theoretical stability and reactivity criteria for metacommunities with different types of interactions and dispersal rates. These theoretical results also apply to isolated communities ($m = 1$ and $d = 0$), and the corresponding criteria are consistent with established stability and reactivity conditions for single isolated communities [32]. The parameters are: $\sigma_1^2 = \sigma^2 - \mathbb{E}^2(|z|)$, $\sigma_2^2 = \sigma^2 - c\mathbb{E}^2(|z|)$, $\sigma_3^2 = \sigma^2 + (1-2c)\mathbb{E}^2(|z|)$, where $m' = m - 1$ and $z$ represents the value taken by the interaction $a_{ij}^k$.

Importantly, these three dynamical regimes have distinct implications for species persistence under ecological disturbances. To investigate this, we model ecological interactions within each patch using a generalized Lotka–Volterra framework. As illustrated schematically in Fig. 1c,d, species persist in stable metacommunities because perturbations decay (Fig. 1c), whereas reactive metacommunities can experience species extinctions under sufficiently frequent perturbations as transient perturbations are amplified. In unstable metacommunities, by contrast, even a single perturbation can trigger extinctions (Fig. 1c). We test these predictions numerically below and quantify the probability of global extinction across randomly sampled metacommunities. In our model, global and local extinctions coincide because species are maintained at low densities through dispersal between patches; in ecological systems, these low-density states would instead correspond to local extinctions.

### Dispersal stabilizes metacommunities

To test our theory, we first compare its predictions with established findings on well-mixed, spatially unstructured communities. Previous studies have shown that stability is governed by a single effective parameter – ecological complexity – which increases with the connectivity of ecological networks or the number of species (Fig. 2a) [11–13, 17–20]. Critically, prior work showed that, as complexity increases, the communities become reactive [32] and subsequently unstable [11], suggesting a universal pathway to instability. Our work extends these conclusions to metacommunities, revealing that diversity and complexity drive a sequential transition from stability to reactivity and eventually to instability (*d* →0 limit illustrated in Fig. 2b,c and Supplementary Figs. 1-3). Similarly, previous studies emphasized that competitive (− */* −) rather than mutualistic (+*/*+) interactions stabilize well-mixed communities — a pattern mirrored in our metacommunity analysis [12, 13, 34], where an increasing proportion of mutualistic interactions has a destabilizing effect (Fig. 2d). For completeness, this analysis can be extended to predatorprey interactions (+*/*−), that have a non-monotonic effect on stability: when competitive interactions dominate, predator-prey dynamics enhance stability; conversely, when mutualistic interactions prevail, excessive predator-prey interactions destabilize the system (Supplementary Fig. 4). The fact that we can reproduce previous results for well-mixed systems validates our framework. Since this new framework additionally incorporates spatial dynamics, we can also explore the dynamics beyond well-mixed conditions. Specifically, we further investigate how dispersal alters system behavior. Importantly, we find that dispersal enhances stability, postponing the onset of reactivity and instability to higher levels of complexity (Fig. 2b,c). This means that dispersal stabilizes metacommunities with stronger ecological interactions and higher diversity that could not be stable in the absence of dispersal. Furthermore, dispersal alleviates the constraint imposed by mutualistic interactions on community stability, enabling metacommunities to remain stable even under a high proportion of mutualistic interactions (Fig. 2d). The strength of this stabilizing effect depends on the dispersal rate *d*: higher dispersal rates enable metacommunities to remain stable at greater levels of complexity, eventually saturating at large values of *d*. This suggests that we can restore stability and prevent systemic collapse by conserving spatial dispersal among different habitats.

**FIG. 2.**
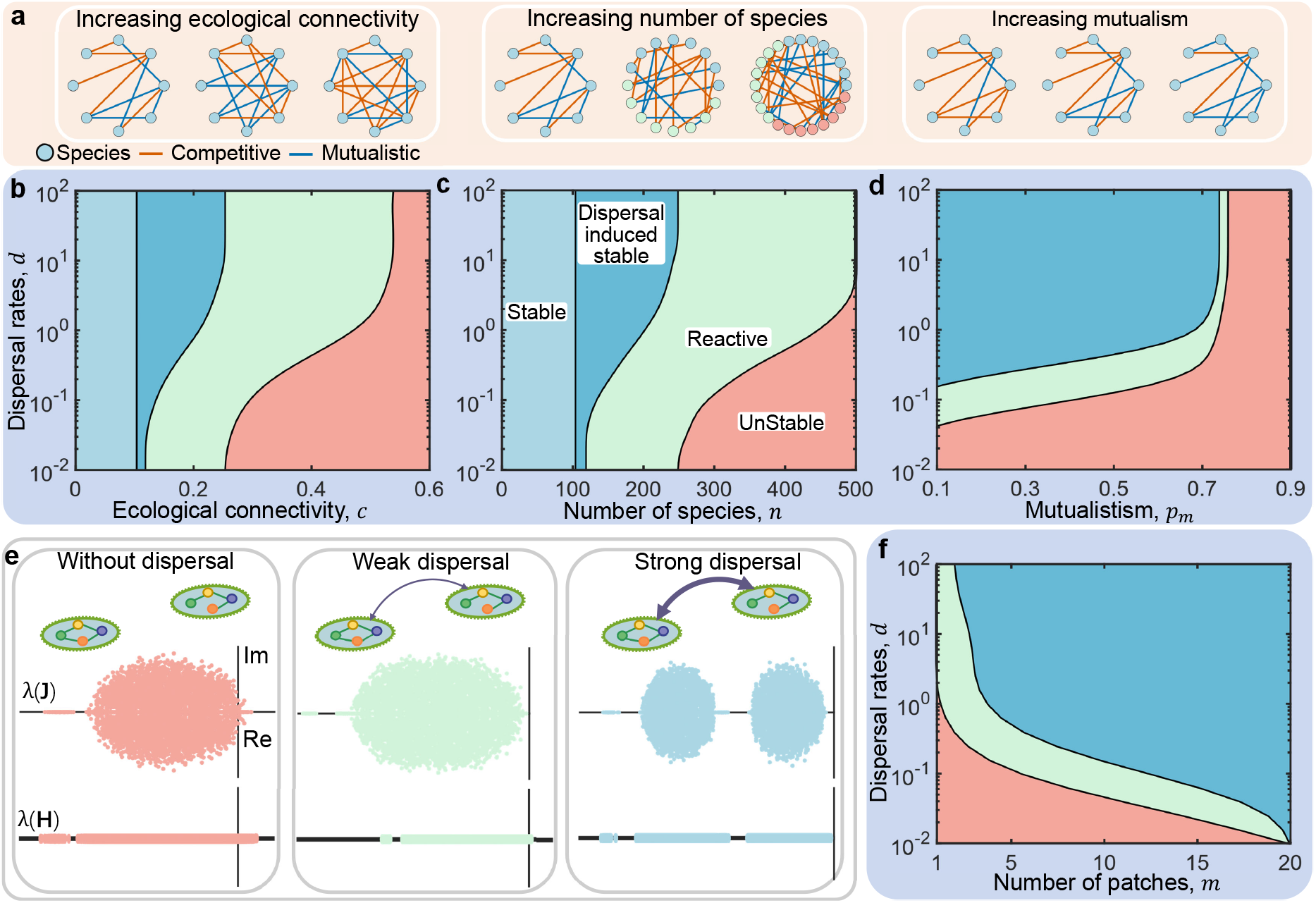
Dispersal between patches is key for the stability of metacommunities. **a**. Structural complexity increases with connectivity, species diversity, and mixed competitive–mutualistic interactions. Orange lines denote competition; blue lines denote mutualism. **b**. Increasing connectivity drives the system toward instability, while dispersal promotes stability by enlarging the early-warning reactive region and shrinking the unstable region. Metacommunities transition from stable (blue) to reactive (green) and then to unstable (red) states. Dark-blue regions are reactive or unstable without dispersal but are stabilized when dispersal is present. **c-d**. Higher species diversity (*m*, **c**) and greater proportion of mutualistic interactions (*p*_*m*_, **d**) reduce inherent stability, but enhanced dispersal buffers these effects, promoting stability. **e**. Mechanisms of dispersal-induced stability: increasing dispersal shifts the eigenvalue spectra of the Jacobian **J** and its symmetric part **H** leftward, turning max ℜ(*λ*(**J**)) and *λ*_max_(**H**) from positive to negative, driving the system from instability to stability. **f**. Greater habitat availability lowers the dispersal threshold required for stability at a given structural complexity. Parameters: *σ* = 0.2, *s* = 1, *n* = 100 (**b-f**), *m* = 2 (**b-e**), *c* = 0.1 (**c-e**) or *c* = 0.4 (**f**).

Mathematically, the stabilizing effect of dispersal can be understood from the eigenvalue spectra, which measure the degree of destabilization in a metacommunity: max ℜ(*λ*(**J**)) *>* 0 indicates an unstable system and larger max ℜ(*λ*(**J**)) signifies greater instability (Fig. 2e). In the absence of dispersal, the metacommunity degenerates into two isolated, unstable communities, given by two independent spectra (blue in Fig. 2e). As dispersal increases, the eigenvalue spectra of **J** and **H** shift leftward (green and orange in Fig. 2e), causing the system to become reactive and stable as the dispersal becomes strong.

In summary, we demonstrate that dispersal can stabilize ecologically complex metacommunities. Importantly, it is not only the dispersal rate but also the number of patches involved in dispersal that jointly determine this effect of dispersal-induced stability (Fig. 2f). Sufficient dispersal between a sufficiently large number of patches allows metacommunities to be stable. However, as either dispersal rate or patch number declines, metacommunities transition to reactive and ultimately unstable states. While a greater number of patches can partially compensate for weak dispersal and *vice versa*, both the number of patches and dispersal rate must be sufficiently large to maintain a truly stable metacommunity.

### Dispersal heterogeneity promotes reactivity

In the previous section, we showed that dispersal is key for the stability of metacommunities. However, these predictions were based on our analytics in Table 1, which rest on the assumption that dispersal is homogeneous. In practice, however, dispersal need not be homogeneous and can vary across scales and in magnitude. In terms of scale, dispersal can either be limited to neighbouring patches (short-range dispersal) [35, 36] or it can be maintained across longer distances (long-range dispersal) [37, 38]. Indeed, certain species, such as birds and fish, are capable of seasonal long-distance migration, and plant seeds can also be dispersed over long distances via birds and other vectors [39, 40], and such long-range dispersal is more frequent with anthropogenic activity [37, 38]. For this reason, we will model both short-range dispersal, either as homogeneous (Fig. 3a) or as heterogeneous (Fig. 3b) in magnitude, as well as long-range dispersal (Fig. 3c).

**FIG. 3.**
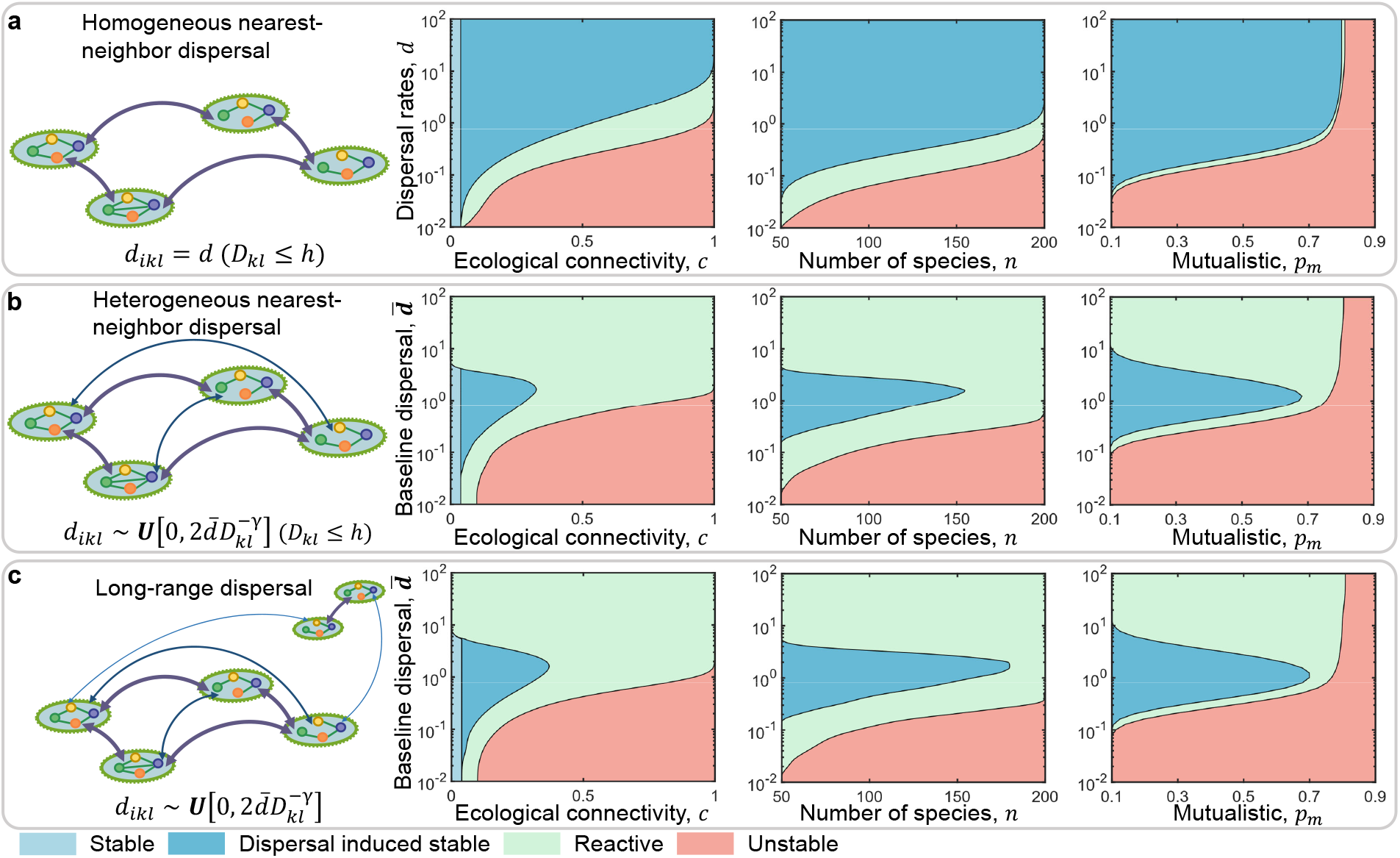
Dispersal-induced stability saturates under homogeneous dispersal but peaks at intermediate intensity under heterogeneous dispersal. **a**. Homogeneous nearest-neighbour dispersal promotes stability, but its stabilizing effect saturates once the dispersal rate exceeds a critical threshold. Increasing dispersal mitigates instability driven by connectivity (left), species diversity (middle), and mutualistic interactions (right) in a 20-patch metacommunity with 2*h* nearest-neighbour dispersal. **b**. Heterogeneous nearest-neighbour dispersal maximizes stability at intermediate intensity, whereas strong dispersal propagates fluctuations across patches, amplifying reactivity. For patches separated by a topological distance *D*_*kl*_ ≤ *h*, dispersal rates satisfy 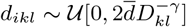, where 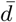 is the baseline dispersal rate. **c**. Heterogeneous long-range dispersal further stabilizes metacommunities. Random addition of 20 long-range dispersal links between patches with *D*_*kl*_ *> h* enhances stability. Across all panels, increasing connectivity, species diversity, and mutualistic interactions destabilize the system, while dispersal modulates this effect. The other parameters are *n* = 50, *s* = 1, *c* = 0.1, *h* = 3, *γ* = 1, and *σ* = 0.4.

We first consider homogeneous nearest-neighbour dispersal, defined by an identical dispersal rate *d* between any two connected patches, as illustrated in Fig. 3a. For simplicity, we imagine that the patches are positioned along a circle, with each patch connected to its 2*h* nearest patches along the circle. Mathematically, the topological distance *D*_*kl*_ can be defined as *D*_*kl*_ = min {|*k* −*l*|, *m*− |*k*− *l*|}. For sufficiently large *h*, we recover the fully connected network that we studied before. For *h* = 1, the patches are connected in a circular topology, providing us with the nearest-neighbour dispersal studied here. Similar to the full-dispersal scenario shown in Fig. 2b–d, increasing complexity drives the metacommunity from stability toward instability. Again, increasing dispersal rate reduces reactivity and instability, while promoting stability. Hence, maintaining spatial dispersal among patches is crucial for sustaining metacommunity stability, where we note that this effect saturates at high dispersal rates.

While the results are similar between the fully connected and sparsely connected networks of patches, we have still ignored the fact that dispersal can be heterogeneous in its magnitude. In metacommunities, dispersal heterogeneity is frequently distance-dependent, with dispersal rates decreasing as spatial separation increases [41, 42]. Accordingly, we assume that dispersal decays with topological distance following a power law with exponent *γ >* 0 and we also incorporate species-level heterogeneity. In this study, we assume that the dispersal rate of species *i* from patch *k* that is connected to patch *l* (*d*_*ikl*_) is drawn from a uniform distribution,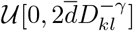. Similar to the case of homogeneous dispersal, we observe that increasing dispersal shifts the system toward stability at higher levels of complexity, where the stable regime seems to dominate at intermediate values of *d* (Fig. 3b). However, under strong dispersal, heterogeneous metacommunities fail to maintain true stability at high complexity levels as homogeneous metacommunities do, and instead exhibit reactivity, even at a low level of ecological complexity (such as when species interact weakly within patches, or their diversity is low). In particular, even intrinsically stable isolated local communities become reactive following coupling by strong heterogeneous dispersal. Intuitively, moderate dispersal allows populations to track favorable conditions across patches, damping local perturbations. Strong dispersal, however, rapidly transmits fluctuations through heterogeneous dispersal, sustaining coexistence but amplifying transient responses [43–45]. Thus, heterogeneous, distance-dependent dispersal most effectively stabilizes metacommunities at intermediate rates, balancing local buffering and system-wide excitation. In the Supplementary Information, we further show that this strong dispersal-induced reactivity disappears if heterogeneity is determined solely by interpatch distance and is the same for each species 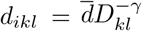; Supplementary Figs. 5 and 6), demonstrating that dispersal heterogeneity between different species is the key factor in generating this dispersal-induced reactivity.

Finally, dispersal might not be limited to nearest patches but may occur across long distances. Here, we assess the effect of long-range dispersal on ecosystem stability by introducing sparse random long-range links into our *h*-nearest-neighbour metacommunities, such that long-range dispersal remains much less frequent than short-distance dispersal. Figure 3c shows the change in stability transition with respect to the baseline dispersal rate after randomly introducing 20 long-range dispersal links into heterogeneous nearest-neighbour metacommunities. Crucially, introduction of non-local dispersal does not have any qualitative effect. Again, consistent with Fig. 3b, stability exhibits a non-monotonic dependence on dispersal rate, reaching a maximum at intermediate values. While long-range dispersal does not qualitatively alter this pattern, it moderately enhances metacommunity stability and expands the reactivity region, which suggests that both long-range and short-range dispersal are key for the stability of metacommunities.

Overall, we extend the study of stability and reactivity in metacommunities to scenarios with heterogeneous dispersal. We find that the stability of metacommunities with heterogeneous dispersal is maximized at intermediate dispersal rates, and reveal that the origin of reactivity in strongly dispersing metacommunities lies in species-specific differences in dispersal. While long-range dispersal moderately stabilizes metacommunities with heterogeneous dispersal, it cannot counteract the rise in system reactivity induced by strong heterogeneous dispersal.

#### From ecological theory to biodiversity conservation

We have analyzed how the stability of metacommunities depends on the spatial structure of dispersal between isolated communities (Figs. 2 and 3). Having established the link between dispersal and stability, we can return to our initial question: how does habitat degradation influence the stability of metacommunities?

Indeed, this question is of primary concern. The data from [2] in Fig. 4a reveals that habitat degradation represents the primary threat to biodiversity across different taxa (amphibians, birds, fish, mammals and reptiles) and regions (North America, South America, Europe, Africa, and Asia). These data demonstrate that, on average, habitat degradation contributes to nearly half of global biodiversity decline. It is therefore critical to assess how this major threat affects the stability of metacommunities in the face of stochastic perturbations, such as pollution incidents, disease outbreaks, or floods and wildfires that are intensified under climate change [8].

**FIG. 4.**
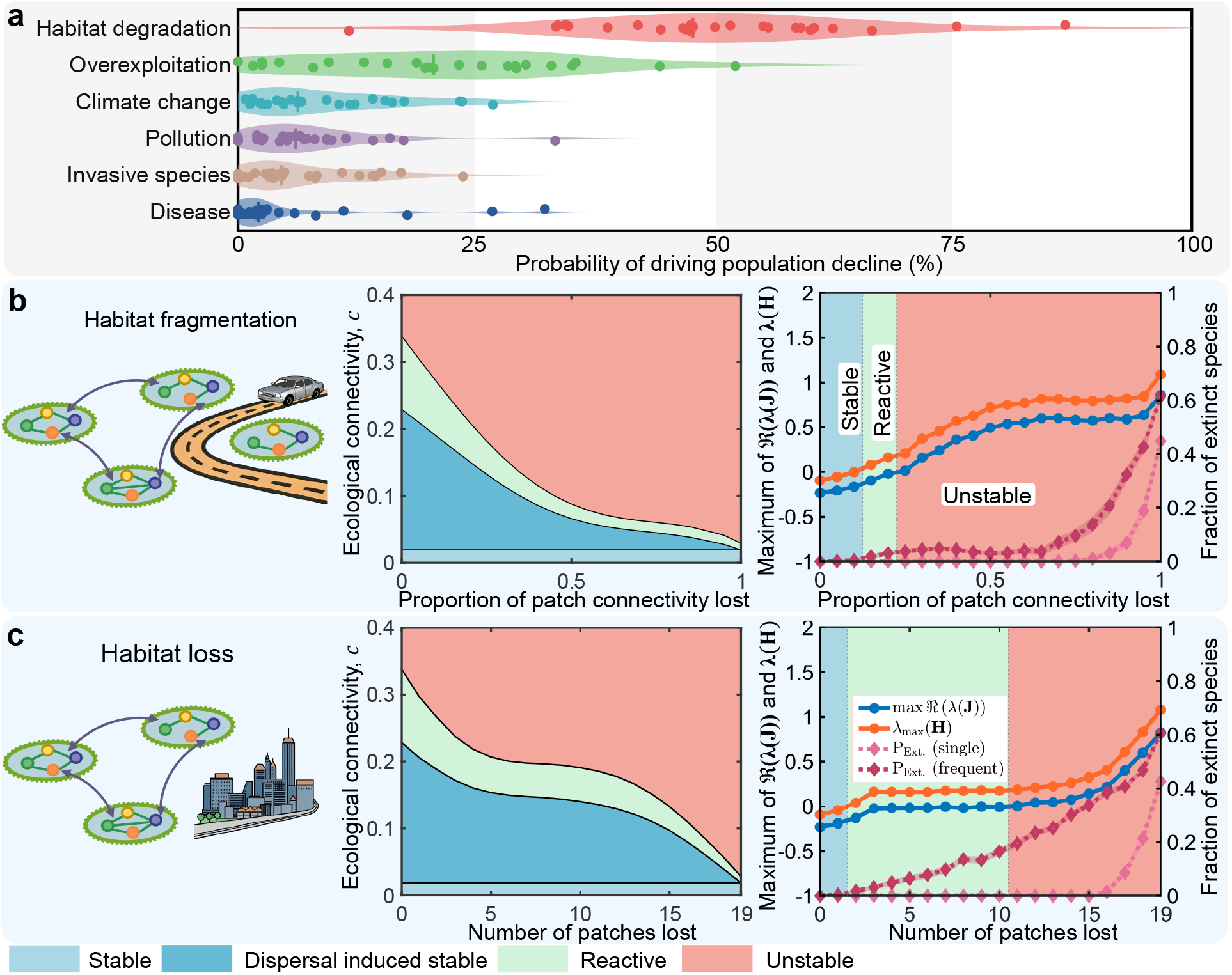
Habitat degradation destabilizes metacommunities. **a**. The probability that a given threat drives population decline. Each point corresponds to a taxon of species in a specific global geographical region, as reported in [2]. **b**. Habitat fragmentation reduces dispersal and drives competitive metacommunities toward instability. Progressive patch connectivity loss can trigger catastrophic collapses of metacommunity stability. **c**. Loss of even a small amount of habitat severely disrupts competitive metacommunity stability. The right panels of **b** and **c** link habitat degradation-induced stability transitions to species extinction. They show the maximum real part of the Jacobian eigenvalues, max ℜ(*λ*(**J**)) (blue), the leading eigenvalue of its symmetric part, *λ*_max_(**H**) (orange), and the mean fraction of extinct species (*P*_Ext._, right axis) following a single perturbation (light red) or repeated perturbations (30 events separated by one time unit; dark red). Shaded areas denote the 95 % confidence intervals based on 50 simulation replicates. In the stable regime, species persist following both single and repeated perturbations, with extinction events rarely observed across stochastic simulations. In the reactive regime, species persist after isolated perturbations, whereas repeated perturbations induce species loss through transient amplification, with extinctions emerging after 30 perturbation events under both patch-connectivity loss (**b**) and patch loss (**c**). Once the metacommunity becomes unstable, even a single perturbation can trigger species extinctions, which become more pronounced under repeated perturbations. These results establish a direct link between stability transitions and population persistence, while showing that reactivity represents not only an early warning of impending instability but also a regime of elevated extinction risk under recurrent disturbances. Parameters are *m* = 20, *n* = 50, *σ* = 0.4, *s* = 1, *h* = 3, *d* = 0.5, and *c* = 0.2.

To explore the impact of habitat degradation on ecological stability, we decompose it into two distinct components: habitat fragmentation and habitat loss. Habitat fragmentation is defined as the reduction of patch connectivity, such as through the construction of roads [46, 47], whereas habitat loss is defined as the loss of habitable patches, such as by turning wild habitats into agricultural or urban land [48, 49]. To apply our theoretical framework, we evaluate the effects of habitat fragmentation and habitat loss separately.

Our new results demonstrate that dispersal among many patches is crucial for maintaining the stability of metacommunities (Figs. 2 and 3). Since habitat fragmentation disrupts the spatial dispersal between patches, our theory predicts that ecological stability will decline as the connectivity of the dispersal network is destroyed (Fig. 4b). Our results reveal a triphasic response. In the initial phase, stability and reactivity decline sharply as dispersal loss weakens connectivity but does not isolate patches. During the intermediate phase, the partially disconnected network maintains a relatively stable core, leaving stability largely unchanged. In the final phase, complete fragmentation isolates communities, triggering an abrupt collapse of stability and contraction of the reactive region.

By the same logic, our study predicts a decline in stability with habitat loss, that is, as patches are lost (Fig. 4c). This triphasic response reflects the changing role of dispersal: in initial phase, disrupts dispersal pathways, weakening spatial coupling and reducing stability; in the intermediate phase, the remaining network retains sufficient connectivity, buffering against further deterioration; in the final stage, continued patch removal diminishes average dispersal capacity, progressively destabilizing the system.

Our theoretical framework for ecological stability can be directly linked to biodiversity persistence by quantifying global species extinctions using generalized Lotka–Volterra metacommunity dynamics (Fig. 4b,c and Supplementary Figs. 7 and 8). Both habitat fragmentation and habitat loss drive metacommunities from stable to reactive and ultimately to unstable regimes, with markedly different consequences for population persistence under different disturbance scenarios. In the stable regime, species persist under both isolated and moderately repeated perturbations, because perturbations decay without generating sufficient transient amplification. In the reactive regime, species persist following a single perturbation but extinctions emerge under sufficiently frequent perturbations, whereas in the unstable regime even a single perturbation can trigger species loss, with repeated perturbations further increasing extinction risk. These results establish a direct connection between linear stability theory and population persistence, showing how habitat degradation progressively increases extinction risk by driving the reduction in metacommunity stability. Notably, asymptotic stability alone does not fully characterize population persistence under recurrent disturbances: systems that remain asymptotically stable may nevertheless experience extinction once they enter the reactive regime, where transient amplification prevents recovery between successive disturbances. Importantly, the emergence of reactivity can precede substantial biodiversity loss, providing an early warning that the system has become increasingly vulnerable to extinction under recurrent disturbances.

## Discussion

Overall, we develop a general theory for the stability of metacommunities, spatially extended communities connected by dispersal. This framework builds on May’s foundational analysis of stability in isolated, well-mixed communities [11, 33], and extends prior efforts to incorporate spatial dispersal [23, 24] in two key ways. First, we allow for a broad spectrum of ecological interactions (Table 1), moving beyond the neutral interactions in [23] and the predator–prey interactions in [24]. Second, we explicitly examine how spatial dispersal shapes reactivity (Fig. 1 and Table 1) – the transient amplification of perturbations. Reactivity has been widely studied in local, well-mixed systems [28–31] and has recently been linked to elevated extinction risk under frequent perturbations [32]. By incorporating general ecological interactions and reactivity, and by linking stability transitions to changes in population persistence, our framework provides a general theory for assessing the impacts of spatial dispersal on extinction risks in metacommunities.

A central result of our theory is that dispersal stabilizes metacommunities (Fig. 2), consistent with previous work [23]. This finding speaks directly to the long-standing debate over the relationship between species diversity and stability. Most theoretical studies, following May [11], predict that increasing diversity reduces stability [13, 50–52]. However, alternative perspectives have questioned the generality of this result [53, 54], noting the apparent stability of highly diverse ecosystems such as rainforests and coral reefs. Our results reconcile these views: although adding more species is destabilizing, dispersal can stabilize systems that would otherwise be unstable (Figs. 2c and 3). In this way, spatial structure buffers against instability, providing a plausible explanation for the persistence of diverse, spatially extended ecosystems. This interpretation is consistent with recent work [55], which shows that dispersal can generate effective sublinear growth, a mechanism linked to the stability of diverse communities [54].

To date, the diversity–stability debate has remained largely theoretical [13, 50–54]. Here, we extend this framework to a central problem in biodiversity conservation: habitat degradation [1–4, 56]. Although habitat degradation is a major driver of population decline (Fig. 4a), current understanding relies largely on field studies and meta-analyses [48, 56–59], which often yield conflicting conclusions and confound causal effects [59]. There is broad agreement that habitat loss drives ecosystem decline beyond a proportional reduction in species abundances [48]. By contrast, the effects of habitat fragmentation *per se* remain debated [59], with some studies reporting strong negative effects on biodiversity [58] and others suggesting weak effects compared to habitat loss [57].

Our model helps resolve this fundamental problem in biodiversity conservation. In line with the generally accepted empirical evidence [48], it predicts that habitat loss reduces metacommunity stability (Fig. 4c), making communities more likely to reorganize following perturbation. Moreover, habitat fragmentation that reduces dispersal connectivity can be comparably destabilizing to the habitat loss (cf. Fig. 4b,c), supporting the empirical evidence that fragmentation can strongly erode biodiversity [58] and should not be neglected in conservation policies. More broadly, our framework connects stability theory to a practical conservation problem by showing that both habitat loss and fragmentation systematically reduce metacommunity stability and increase extinction risk under recurrent disturbances, even before the system crosses the instability boundary (Figs. 2f and 4b,c). This distinction is important because asymptotically stable metacommunities may remain viable under isolated perturbations yet become vulnerable when perturbations recur faster than ecological recovery. In doing so, it complements previous theoretical approaches that emphasized trophic collapse [60, 61], reduced species persistence [62], or elevated temporal variability [63], but did not focus on recovery following perturbation.

Our findings reveal two complementary strategies for biodiversity conservation: enhancing species dispersal and expanding habitat area. The former can be achieved by conserving and restoring ecological corridors to ensure gene flow [64–66], whereas the latter relies on habitat protection and restoration [67, 68]. In practical conservation, rewilding is far more difficult and resource-intensive than protecting existing habitats, leading current policies to prioritize habitat conservation. However, the number of species that an ecosystem can sustain is jointly determined by dispersal capacity and the number of habitat patches (Figs. 2f and 4) [56, 69, 70]. If dispersal is improved without sufficient habitat capacity, biodiversity cannot be effectively maintained. Thus, habitat protection and rewilding are not mutually exclusive alternatives; instead, they constitute two indispensable pillars of biodiversity conservation that must be pursued together.

Despite its generality, our framework is restricted to the local stability of metacommunities near feasible steady states. As a result, it relies on small perturbations around equilibrium points and does not capture more complex nonlinear dynamics. Although we partially capture transient dynamics by reactivity, our framework cannot resolve the specific forms these transients may take, including oscillations [71], Turing instabilities [72], and chaos [73, 74]. Extending this framework to capture such transient regimes, building on recent theoretical advances [75], will be an important direction for future research. Furthermore, because local stability is difficult to quantify in empirical systems, further work will be needed to connect it to alternative metrics of stability, including variability [76], vulnerability [77] and resilience [78]. Finally, although habitat degradation is a major driver of population decline (Fig. 4a), we focus here exclusively on this threat and treat other drivers of biodiversity loss as external perturbations. Extending the framework to these additional threats remains an important direction for future research. In the Supplementary Information, for example, we show that invasive species can further destabilize metacommunities (Supplementary Fig. 9).

Overall, we establish a general theoretical framework for the stability of metacommunities and show its applicability to open problems beyond theoretical ecology. In particular, we apply this framework to a central question in biodiversity conservation, showing that both habitat loss and fragmentation drive metacommunities across stability boundaries, ultimately reducing biodiversity persistence. More broadly, our results extend May’s stability theory from spatially well-mixed communities towards practical conservation problems in increasingly fragmented and disturbance-prone ecosystems.

## Methods

### Estimation of the eigenvalues of the Jacobian matrix

In contrast to single community analyses, obtaining the eigenvalue spectra of the Jacobian matrix **J** and its symmetric counterpart **H** in metacommunities is challenging, as they vary continuously with dispersal rate. We therefore estimate the largest real parts of the eigenvalues of **J** and **H** under weak and strong dispersal to derive analytical stability and reactivity criteria, which relate system behavior to the key ecological parameters of complexity (*n, m, c, σ*) and the dispersal rate (*d*). For large metacommunities, low rank perturbations of large random matrices yield approximations of max ℜ(*λ*(**J**)) and *λ*_max_(**H**) under weak dispersal (small *d*) [79–81], giving max

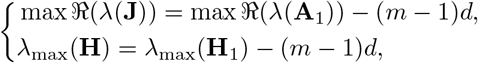

in which 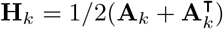. Conversely, under strong dispersal (large *d*), by employing perturbation theory of eigenvalues [82], we derive approximations for eigenvalues with the largest real parts of **J** and **H** as

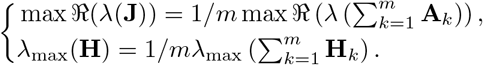

Further details are provided in the Supplementary Information.

### Constructing the interaction matrix with mixed interactions

Mixed interactions imply that species interact through three modes—predator-prey, competition, and mutualism-each occurring with specific probabilities: *p*_+*/−*_, *p*_*−/−*_, and *p*_+*/*+_. The construction algorithm for a metacommunity with mixed interactions is as follows: i) Initialize the interaction matrix *A*_*k*_ with zeros; ii) Set all diagonal terms to −*s* (i.e., 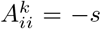); iii) For each off-diagonal pair 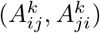 where *i < j*, draw a random number *x* from a uniform distribution *U*[0, 1]; iv) If *x* ≥ *c*, assign zero to both 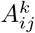 and 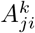; v) If *x < c*, draw another random number *y* from *U*[0, 1]; vi) If *y* ≤ *p*_+*/−*_, with equal probability draw either 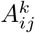 from the negative half-normal distribution −|*N*(0, *σ*^2^)| and 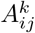 from the half-normal distribution ~|*N* (0, *σ*^2^)|, or the opposite (predator-prey interaction); vii) If *p*_+*/−*_ *< y* ≤ *p*_+*/−*_ + *p*_*−/−*_, draw 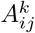 from the negative half-normal distribution −|*N*(0, *σ*^2^)| and 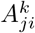 from the negative half-normal distribution −|*N*(0, *σ*^2^)|; otherwise, draw 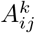 from the positive half-normal distribution |*N* (0, *σ*^2^)| and 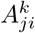 from the positive half-normal distribution |*N* (0, *σ*^2^)|.

The statistical properties of the interaction matrix **A**_*k*_ with mixed interaction can be extracted

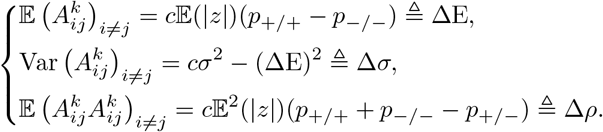

### Stability and reactivity criteria of mixed interaction metacommunities

Based on these statistics, we have 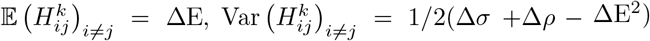, and 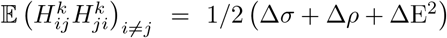. According to random matrix theory and the low-rank perturbation theorem [79–82], the maximum eigenvalues of the matrices **A**_*k*_ and **H**_*k*_ are

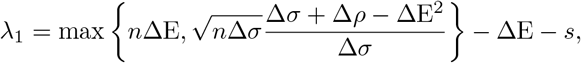

and

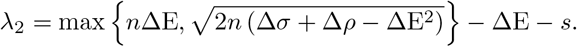

Therefore, under weak dispersal, the stability and reactivity criteria for metacommunities with mixed interactions are,

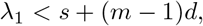

and

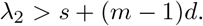

While, under strong dispersal, the stability and reactivity criteria for metacommunities with mixed interactions are

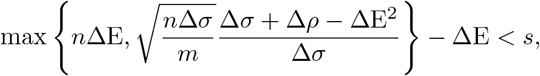

See Supplementary Information for detailed derivation.

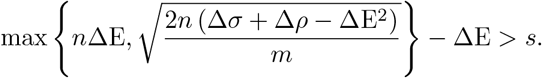

### Constructing the heterogeneous spatial dispersal matrix

For metacommunities with heterogeneous dispersal, we initially examined metacommunities with *h*-nearest-neighbour dispersal, and subsequently considered introducing long-range dispersal into these systems. The interaction matrix **A**_*k*_ is constructed as described previously. The construction algorithm for the dispersal matrices **D**_*kl*_ of metacommunities with *h*-nearest-neighbour dispersal is as follows: i) For each matrix **D**_*kl*_ where *k < l*, if |*l* − *k*| ≤ *h* or |*l* – *m* − *k* | ≤ *h*, assign both **D**_*kl*_ and **D**_*lk*_ to diagonal matrices whose diagonal entries are independently drawn from a uniform distribution 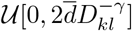. ii) Otherwise, set **D**_*kl*_ = **0**_*n×n*_.

The algorithm for adding long-range dispersal to meta-communities with *h*-nearest-neighbour dispersal is: i) Construct all *h*-nearest-neighbour dispersal matrices **D**_*kl*_ as described above. ii) For every pair **D**_*kl*_ where *k < l*, if |*l* − *k*| *> h* and *m* − |*l* − *k* |*> h*, draw a random number *x* from a uniform distribution *U*[0, 1]. iii) Sort all values of *x*_*kl*_ in descending order, select the top 20 largest entries, and store their corresponding indices (*k, l*) within the index set ℐ. iv) If (*k, l*) ∉ *ℐ*, set **D**_*kl*_ = **D**_*lk*_ = **0**_*n×n*_. v) If (*k, l*) ∈ ℐ, set both matrices to be diagonal matrices whose diagonal entries are drawn from the same, 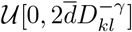.

It should be noted that the uniform distribution, 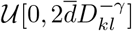 is selected for the simulations; however, other probability distributions are feasible, and the species heterogeneity can be modulated by tuning the variance of the chosen distribution.

## Data availability

All analyzed data are simulated and can be replicated using the provided code.

## Code availability

The codes used for analyses is available at Github (https://github.com/yinxiangxin/Metacommunity.git). May, R. M. Will a large complex system be stable? *Nature* **238**, 413–414 (1972).

## Acknowledgements

The authors thank Professor Kevin R. Foster for his valuable discussions and comments on this work. This work was supported by the National Natural Science Foundation of China (Grant Nos. 12271519, 119251023, 22120102001), the Fundamental Research Funds for the Central Universities (Grant Nos. 2024KYJD2002 and 2023ZDYQ11005). For the purpose of open access, the author has applied a CC BY public copyright licence to any author accepted manuscript arising from this submission.

## Author contributions

Y.S., X.Y. and P.K.M. initiated and conceived the study. Y.S., X.Y. Z.J. and S.B. performed theoretical analysis and numerical simulations. Y.S., V.P., W.L. and P.K.M designed the study and analyzed the results. Y.S., X.Y. and V.P. wrote the first draft of the manuscript, and all authors edited the manuscript. Y.S. and X.Y. contributed equally to this work.

## Competing interests

The authors declare no competing interests.

## Additional information

Supplementary Information is available for this paper.

## Notes

https://github.com/yinxiangxin/Metacommunity.git

### Competing Interest Statement

The authors have declared no competing interest.

## References

[1] Díaz, S. et al. Pervasive human-driven decline of life on earth points to the need for transformative change. Science 366, eaax3100 (2019).

[2] World Wildlife Fund. 2024 living planet report (2024). URL https://www.worldwildlife.org/publications/2024-living-planet-report/.

[3] Armstrong McKay, D. I. et al. Exceeding 1.5°c global warming could trigger multiple climate tipping points. Science 377, eabn7950 (2022).

[4] Li, G., Wang, Z., Sun, S., Qi, W. & Watson, J. E. Projected human land-use pressures and natural habitat conversion risk within global terrestrial protected areas. Nat. Ecol. Evol. 10, 281–292 (2026).

[5] Fahrig, L. Effects of habitat fragmentation on biodiversity. Annu. Rev. Ecol. Evol. Syst. 34, 487–515 (2003).

[6] Morales, H. E. & Cock van Oosterhout, C. The impact of habitat loss and population fragmentation on genomic erosion. Conserv. Genet. 25, 49–57 (2024).

[7] Pearse, I. S., Baty, J. H., Pesendorfer, M. B., Olson, R. & Koenig, W. D. Oak masting breaks down in a highly fragmented, pollen-limited landscape. Proc. Natl. Acad. Sci. U. S. A. 123, e2533900123 (2026).

[8] Masson-Delmotte, V. et al. Climate change 2021: the physical science basis. Contribution of working group I to the sixth assessment report of the intergovernmental panel on climate change 2, 2391 (2021).

[9] Xu, C., Kohler, T. A., Lenton, T. M.Svenning, J.-C. & Scheffer, M. Future of the human climate niche. Proc. Natl. Acad. Sci. U. S. A. 117, 11350–11355 (2020).

[10] Shurety, A. L., Thompson, M. S., Couce, E., Cameron, T. C. & O’Gorman, E. J. Commercial fishing amplifies impacts of increasing temperature on predator-prey interactions in marine ecosystems. Nat. Commun. 17, 628 (2026).

[11] May, R. M. Will a large complex system be stable? Nature 238, 413–414 (1972).

[12] Mougi, A. & Kondoh, M. Diversity of interaction types and ecological community stability. Science 337, 349–351 (2012).

[13] Coyte, K. Z., Schluter, J. & Foster, K. R. The ecology of the microbiome: networks, competition, and stability. Science 350, 663–666 (2015).

[14] Azaele, S. & Maritan, A. Generalized dynamical mean field theory for non-gaussian interactions. Phys. Rev. Lett. 133, 127401 (2024).

[15] Yang, Y., Foster, K. R., Coyte, K. Z. & Li, A. Time delays modulate the stability of complex ecosystems. Nat. Ecol. Evol. 7, 1610–1619 (2023).

[16] Qin, B.-W. et al. Reordered hierarchical complexity in ecosystems with delayed interactions. PNAS Nexus 4, pgaf214 (2025).

[17] Srednick, G. & Swearer, S. E. Understanding diversity–synchrony–stability relationships in multitrophic communities. Nat. Ecol. Evol. 8, 1259–1269 (2024).

[18] Gilarranz, L. J., Rayfield, B., Liñán-Cembrano, G., Bascompte, J. & Gonzalez, A. Effects of network modularity on the spread of perturbation impact in experimental metapopulations. Science 357, 199–201 (2017).

[19] Grilli, J., Rogers, T. & Allesina, S. Modularity and stability in ecological communities. Nat. Commun. 7, 12031 (2016).

[20] Grilli, J., Barabás, G., Michalska-Smith, M. J. & Allesina, S. Higher-order interactions stabilize dynamics in competitive network models. Nature 548, 210–213 (2017).

[21] Tao, Y., Hastings, A., Lafferty, K. D., Hanski, I. & Ovaskainen, O. Landscape fragmentation overturns classical metapopulation thinking. Proc. Natl Acad. Sci. U. S. A. 121, e2303846121 (2024).

[22] Carrara, F., Altermatt, F., Rodriguez-Iturbe, I. & Rinaldo, A. Dendritic connectivity controls biodiversity patterns in experimental metacommunities. Proc. Natl Acad. Sci. U. S. A. 109, 5761–5766 (2012).

[23] Gravel, D., Massol, F. & Leibold, M. A. Stability and complexity in model meta-ecosystems. Nat. Commun. 7, 12457 (2016).

[24] Baron, J. W. & Galla, T. Dispersal-induced instability in complex ecosystems. Nat. Commun. 11, 6032 (2020).

[25] Hastings, A., McCann, K. & Gellner, G. Competition–colonization trade-off can explain any observed abundances and assumed competitive hierarchies. Proc. Natl. Acad. Sci. U. S. A. 122, e2526827122 (2025).

[26] Jackson, Z., Leibold, M. A., Holt, R. D. & Xue, B. Modeling and inferring metacommunity dynamics with maximum caliber. Proc. Natl. Acad. Sci. U. S. A. 123, e2520867123 (2026).

[27] Neubert, M. G. & Caswell, H. Alternatives to resilience for measuring the responses of ecological systems to perturbations. Ecology 78, 653–665 (1997).

[28] Tang, S. & Allesina, S. Reactivity and stability of large ecosystems. Front. Ecol. Evol. 2, 21 (2014).

[29] Suweis, S., Grilli, J., Banavar, J. R., Allesina, S. & Maritan, Effect of localization on the stability of mutualistic ecological networks. Nat. Commun. 6, 10179 (2015).

[30] Suweis, S., Carr, J. A., Maritan, A., Rinaldo, A. & D’Odorico, P. Resilience and reactivity of global food security. Proc. Natl. Acad. Sci. U. S. A. 112, 6902–6907 (2015).

[31] Fruchart, M., Hanai, R., Littlewood, P. B. & Vitelli, V. Non-reciprocal phase transitions. Nature 592, 363–369 (2021).

[32] Yang, Y., Coyte, K. Z., Foster, K. R. & Li, A. Reactivity of complex communities can be more important than stability. Nat. Commun. 14, 7204 (2023).

[33] May, R. M. Stability and Complexity in Model Ecosystems (Princeton University Press, Princeton, 1973).

[34] Palmer, J. D. & Foster, K. R. Bacterial species rarely work together. Science 376, 581–582 (2022).

[35] Wu, N. C. & Seebacher, F. Physiology can predict animal activity, exploration, and dispersal. Commun. Biol. 5, 109 (2022).

[36] Lamers, K. P., Nilsson, J.-Å., Nicolaus, M. & Both, C. Adaptation to climate change through dispersal and inherited timing in an avian migrant. Nat. Ecol. Evol. 7, 1869–1877 (2023).

[37] Decout, S., Manel, S., Miaud, C. & Luque, S. Integrative approach for landscape-based graph connectivity analysis: a case study with the common frog (rana temporaria) in human-dominated landscapes. Landsc. Ecol. 27, 267–279 (2012).

[38] Nobre, D. A., Abbott, K. C., Machta, J. & Hastings, A. Long-range dispersal promotes spatial synchrony but reduces the length and time scales of synchronous fluctuations. Theor. Ecol. 19, 5 (2026).

[39] González-Varo, J. P. et al. Limited potential for bird migration to disperse plants to cooler latitudes. Nature 595, 75–79 (2021).

[40] Levey, D. J., Bolker, B. M., Tewksbury, J. J., Sargent, S. & Haddad, N. M. Effects of landscape corridors on seed dispersal by birds. Science 309, 146–148 (2005).

[41] Clobert, J., Baguette, M., Benton, T. G. & Bullock, J. M. Dispersal ecology and evolution (Oxford University Press, 2012).

[42] Skelsey, P., With, K. A. & Garrett, K. A. Why dispersal should be maximized at intermediate scales of heterogeneity. Theor. Ecol. 6, 203–211 (2013).

[43] Ellner, S. P. et al. Habitat structure and population persistence in an experimental community. Nature 412, 538–543 (2001).

[44] Dey, S. & Joshi, A. Stability via asynchrony in drosophila metapopulations with low migration rates. Science 312, 434–436 (2006).

[45] Nell, L. A. et al. Dispersal stabilizes coupled ecological and evolutionary dynamics in a host-parasitoid system. Science 383, 1240–1244 (2024).

[46] GonÇalves-Souza, T. et al. Species turnover does not rescue biodiversity in fragmented landscapes. Nature 640, 702–706 (2025).

[47] Ma, F. et al. Biodiversity and habitat complexity buffer the destabilizing effects of anthropogenic activities on riverine fish communities. Nat. Commun. (2026).

[48] Chase, J. M., Blowes, S. A., Knight, T. M., Gerstner, K. & May, F. Ecosystem decay exacerbates biodiversity loss with habitat loss. Nature 584, 238–243 (2020).

[49] Kuzyk, Z. Z. A. et al. Eelgrass ecosystem collapse and social-ecological regime shift driven by hydropower development and climate change. Nat. Commun. 17, 2917 (2026).

[50] Allesina, S. & Tang, S. Stability criteria for complex ecosystems. Nature 483, 205–208 (2012).

[51] Hu, J., Amor, D. R., Barbier, M., Bunin, G. & Gore, J. Emergent phases of ecological diversity and dynamics mapped in microcosms. Science 378, 85–89 (2022).

[52] Yonatan, Y., Amit, G., Friedman, J. & Bashan, A. Complexity–stability trade-off in empirical microbial ecosystems. Nat. Ecol. Evol. 6, 693–700 (2022).

[53] Rozdilsky, I. D. & Stone, L. Complexity can enhance stability in competitive systems. Ecol. Lett. 4, 397–400 (2001).

[54] Hatton, I. A., Mazzarisi, O., Altieri, A. & Smerlak, M. Diversity begets stability: Sublinear growth and competitive coexistence across ecosystems. Science 383, eadg8488 (2024).

[55] Al-Hiyasat, A., Swartz, D. W., Gore, J. & Kardar, M. Spatiotemporal noise stabilizes unbounded diversity in strongly-competitive communities. arXiv preprint arXiv:2602.13423 (2026).

[56] Haddad, N. M. et al. Habitat fragmentation and its lasting impact on earth’s ecosystems. Sci. Adv. 1, e1500052 (2015).

[57] Fahrig, L. Ecological responses to habitat fragmentation per se. Annu. Rev. Ecol. Evol. Syst. 48, 1–23 (2017).

[58] Fletcher, R. J., Jr et al. Is habitat fragmentation good for biodiversity? Biol. Conserv. 226, 9–15 (2018).

[59] Valente, J. J. et al. Toward conciliation in the habitat fragmentation and biodiversity debate. Landsc. Ecol. 38, 2717–2730 (2023).

[60] Solé, R. V. & Montoya, J. M. in Ecological network meltdown from habitat loss and fragmentation 305–323 (Oxford University Press, New York, 2005).

[61] Liao, J., Bearup, D. & Blasius, B. Diverse responses of species to landscape fragmentation in a simple food chain. J. Anim. Ecol. 86, 1169–1178 (2017).

[62] Laurance, W. F. Theory meets reality: How habitat fragmentation research has transcended island biogeographic theory. Biol. Conserv. 141, 1731–1744 (2008).

[63] McWilliams, C., Lurgi, M., Montoya, J. M., Sauve, A. & Montoya, D. The stability of multitrophic communities under habitat loss. Nat. Commun. 10, 2322 (2019).

[64] Damschen, E. I., Haddad, N. M., Orrock, J. L., Tewksbury, J. J. & Levey, D. J. Corridors increase plant species richness at large scales. Science 313, 1284–1286 (2006).

[65] McGuire, J. L., Lawler, J. J., McRae, B. H., Nuñez, T. A. & Theobald, D. M. Achieving climate connectivity in a fragmented landscape. Proc. Natl. Acad. Sci. U. S. A. 113, 7195–7200 (2016).

[66] Naidoo, R. et al. From science to impact: Conserving ecological connectivity in large conservation landscapes. Proc. Natl. Acad. Sci. U. S. A. 122, e2410937122 (2025).

[67] Xu, C. et al. Herbivory limits success of vegetation restoration globally. Science 382, 589–594 (2023).

[68] Langhammer, P. F. et al. The positive impact of conservation action. Science 384, 453–458 (2024).

[69] Benayas, J. M. R., Newton, A. C., Diaz, A. & Bullock, J. M. Enhancement of biodiversity and ecosystem services by ecological restoration: a meta-analysis. Science 325, 1121–1124 (2009).

[70] Strassburg, B. B. et al. Global priority areas for ecosystem restoration. Nature 586, 724–729 (2020).

[71] Hastings, A. et al. Transient phenomena in ecology. Science 361, eaat6412 (2018).

[72] Hajian-Forooshani, Z., Rivera-Salinas, I. S., Perfecto, I. & Vandermeer, J. The population dynamics of clustered consumer–resource spatial patterns: Insights from the demographics of a turing mechanism. Proc. Natl. Acad. Sci. U.S.A. 122, e2407991121 (2025).

[73] Hechler, R. M. & Krkosek, M. Temperature variation and life history mediate nonlinearity in fluctuations of marine fish populations worldwide. Nat. Ecol. Evol. 10, 942–951 (2026).

[74] Lechón-Alonso, P., Kundu, S., Lemos-Costa, P., Capitán, J. A. & Allesina, S. Robust coexistence in competitive ecological communities. Nat. Commun. 17, 2637 (2026).

[75] Yang, Y., Barabás, G., Saavedra, S. & Li, A. Catalysts and inhibitors of critical transitions in ecological systems. Proc. Natl. Acad. Sci. U.S.A. 123, e2516856122 (2026).

[76] Meng, B. et al. Stabilizing effects of biodiversity arise from species-specific dynamics rather than interspecific interactions in grasslands. Nat. Ecol. Evol. 9, 1837–1847 (2025).

[77] Bernardi, D. et al. Dispersal diversity buffers species vulnerability to local extinction. Nat. Ecol. Evol. 1–14 (2026).

[78] Capdevila, P. et al. Life history mediates the trade-offs among different components of demographic resilience. Ecol. Lett. 25, 1566–1579 (2022).

[79] Naumov, A. Elliptic law for real random matrices. arXiv preprint arXiv:1201.1639 (2012).

[80] Nguyen, H. H. & O’Rourke, S. The elliptic law. Int. Math. Res. Notices 2015, 7620–7689 (2015).

[81] O’Rourke, S. & Renfrew, D. Low rank perturbations of large elliptic random matrices. Electron. J. Probab. 19, 1–65 (2014).

[82] Gomez, S. et al. Diffusion dynamics on multiplex networks. Phys. Rev. Lett. 110, 028701 (2013).

